# Metabolic switching of *Mycobacterium tuberculosis* during hypoxia is controlled by the virulence regulator PhoP

**DOI:** 10.1101/845271

**Authors:** Prabhat Ranjan Singh, Vijjamarri Anil Kumar, Dibyendu Sarkar

## Abstract

*Mycobacterium tuberculosis* (Mtb) retains the unique ability to establish an asymptomatic latent infection. A fundamental question in mycobacterial physiology is to understand the mechanisms involved in hypoxic stress, a critical player in persistence. Here, we show that the virulence regulator PhoP responds to hypoxia, the dormancy signal and effectively integrates hypoxia with nitrogen metabolism. We also provide evidence to demonstrate that both under nitrogen limiting conditions and during hypoxia, *phoP* locus controls key genes involved in nitrogen metabolism. Consistently, under hypoxia *ΔphoP* shows growth attenuation even with surplus nitrogen, the alternate electron acceptor, and complementation of the mutant restores bacterial growth. Together, our observations provide new biological insights into the role of PhoP in integrating nitrogen metabolism with hypoxia by the assistance of the hypoxia regulator DosR. The results have significant implications on the mechanism of intracellular survival and growth of the tubercle bacilli under a hypoxic environment within the phagosome.

**Importance:** Mtb retains the unique ability to establish an asymptomatic latent infection. To understand the mechanisms involved in hypoxic stress which plays a critical role in persistence, we show that the virulence regulator PhoP responds to hypoxia, the dormancy signal. In keeping with this, *phoP* was shown to play a major role in Mtb growth under hypoxia even in presence of surplus nitrogen, the alternate electron acceptor. Our results showing regulation of hypoxia-responsive genes provide new biological insights into role of the virulence regulator in metabolic switching by sensing hypoxia and integrating nitrogen metabolism with hypoxia by the assistance of the hypoxia regulator DosR.

## INTRODUCTION

A hallmark of tuberculosis is the unique ability of Mtb to establish an asymptomatic latent infection and persist within granulomas in a dormant form, sometimes for a very long time, before reactivation to cause the active disease. As survival and persistence in this environment depends on the sensing of signals and ability to induce a robust adaptive response, one of the major aspects to understand latent TB relates to understanding molecular mechanism of adaptation of the tubercle bacilli in response to environmental stress.

Despite its requirement of oxygen for growth, during latency Mtb can survive without oxygen for a surprisingly long time. Thus, two *in vivo* conditions are often linked to latent TB. These are hypoxia and exposure to immune effectors such as nitric oxide (NO) (1–3). The ability to produce reactive nitrogen species (RNS) by host-inducible nitric oxide synthase (iNOS) contributes to TB infections by its effect on both the host and the pathogen (4). Therefore, functional iNOS expression could be detected in the lung macrophages of human TB patients (5, 6). Consistently, hypoxia and nitric oxide dependent bacterial adaptation to a dormant state induces latent tuberculosis in mice (1, 3). During limiting oxygen concentrations, Mtb induces reduction of nitrate (NO_3_^−^) to nitrite (NO_2_^−^) to control redox homeostasis and energy production (7). When nitrate is taken up by mycobacteria via passive diffusion (8, 9), the mycobacterial nitrate reductase (encoded by *narGHIJ*) is expressed at a low level under aerobic conditions (7). Also, hypoxia promotes induction of NarK2, a nitrate transporter enabling rapid accumulation of nitrite by Mtb cultures growing under oxygen limiting conditions but in the presence of nitrate (8). Consistent with these results, transcripts from *narG*, encoding a nitrate reductase subunit and *narX*, encoding a non-functional nitrate reductase, were identified within granulomas of human TB samples (10, 11). These results suggest that intracelullar mycobacteria within the human host encounters a very low oxygen tension and most likely adapt to the microenvironment by respiring nitrate. However, the key regulators that connect hypoxia and nitrogen metabolism of mycobacteria, and define the mechanisms that promote and maintain TB latency in humans, remain poorly understood.

Previous studies have shown that upon exposure to hypoxia or nitric oxide, the *dosR-dosS* system activates expression of ~48 genes that are part of the dormancy survival (Dos) regulon (12, 13). While DosR is essential for mycobacterial survival under dormancy (12, 14), phosphorylated DosR induces expression of hypoxia-responsive genes (13). DosR cooperatively binds to target promoters containing a minimum of two tandem binding sites, and the proximal DosR binding site often juxtapose with the −35 sequence element of the promoters (15, 16). In keeping with this, DosR-SigA interaction remains essential in the dormancy survival program of Mtb (17). Although previous studies suggest that Mtb *dosR* is under the regulation of PhoP (18–20) with results supporting PhoP recruitment to *dosR* upstream regulatory region (21–23), role of PhoP during mycobacterial hypoxia remains unknown.

Recently, under anaerobic conditions Δ*dosR* Mtb was shown to be growth defective at low pH (24). Because PhoP controls pH-driven adaptations of Mtb (25–28), in this study we focused to investigate whether virulence-associated *phoP* locus is linked to hypoxic response of the tubercle bacilli. Transcript profiling indicates that induction of *dosR* regulon was subdued in *ΔphoP* mutant, suggesting that PhoP functions as an activator of hypoxia-inducible genes. We provide evidence to show striking increase in nitrate and nitrite reduction under hypoxia (7), a state related to non-replicating persistence of Mtb either for redox balance maintenance or to provide energy during shift-down (8), is controlled by the *phoP* locus. In keeping with these results, *phoP* strongly impacts on the expression of genes related to nitrogen metabolism, and under oxygen austerity even in the presence of surplus nitrogen conditions *ΔphoP* Mtb was significantly growth defective relative to the WT bacilli. Together, these results establish that (a) metabolic switching of Mtb under hypoxia is achieved by integration of hypoxia with nitrogen metabolism, and (b) convergence of PhoP and DosR as co-activators of hypoxia-inducible genes, controlled by protein-protein contacts, coordinate nitrogen metabolism in response to hypoxia.

## RESULTS

### phoP impacts hypoxia-inducible gene expression of Mtb

To investigate whether *phoP* impacts on hypoxia, we compared expression of hypoxia-inducible genes in the WT and *ΔphoP* mutant of Mtb using *in vitro* model of hypoxia (3) (Fig. 1). The results suggest that under normal conditions the major hypoxia-inducible genes, which belong to the *dosR* regulon (12–14), are significantly down-regulated in *ΔphoP* relative to the WT bacilli. However, during hypoxia their expressions remain comparable in the WT and the mutant strain (compare Figs. 1A and 1B). In contrast, *dosR* regulon genes which are involved in N_2_-metabolism (29), shows a *phoP*-dependent activation both during normal conditions and hypoxia (Fig. 1). Because *narG* and *nirB* control reduction of nitrate and nitrite, respectively (29) and hypoxia is accompanied with an increase in nitrate reduction (7), together these results suggest that during hypoxia PhoP appears to activate expression of genes involved in N_2_-metabolism. In keeping with these results, *glnR*, a major regulator of nitrate assimilation (29) is strongly repressed by PhoP during hypoxia but not under normal conditions (compare Figs. 1B and 1A). Although GlnR is known for its role in nitrate assimilation of Mtb (29), repression of *glnR* by PhoP during hypoxia rules out the possibility of activation of *narG* and *nirB* via GlnR. Table S1 lists the PAGE-purified oligonucleotides used in RT-PCR experiments. Although a previous study by Smith and co-workers had identified PhoP-regulated transcriptome, the DNA arrays using exponentially growing cells under normal conditions did not reveal an influence of PhoP on the expression of *dosR* or other hypoxia-inducible genes (30). In contrast, using a different *phoP* mutant strain, Asensio et al showed that PhoP functions as an activator of *dosR* (19). While different experimental conditions and varying quantitative approaches appear to account for this discrepancy, clearly our results are consistent with Asensio et al showing a significant effect of PhoP on the expression of *dosR*. More recently, Vashist et al have shown that PhoP functions as a repressor of *dosR* (23). Although both our study and this report have investigated the regulation of *dosR* by PhoP, use of laboratory attenuated Mtb H37Ra, a different genetic background comprising genomic copy of a mutant PhoP (S219L) by Vasisht et al. versus a clean mutant used in our study-most likely accounts for this discrepancy. Also, it should be noted that except PhoP binding to *nirB* promoter in a genome-wide SELEX experiment (31), till date, there has been no report linking PhoP with hypoxia-inducible genes other than *dosR* itself.

**Fig. 1:**
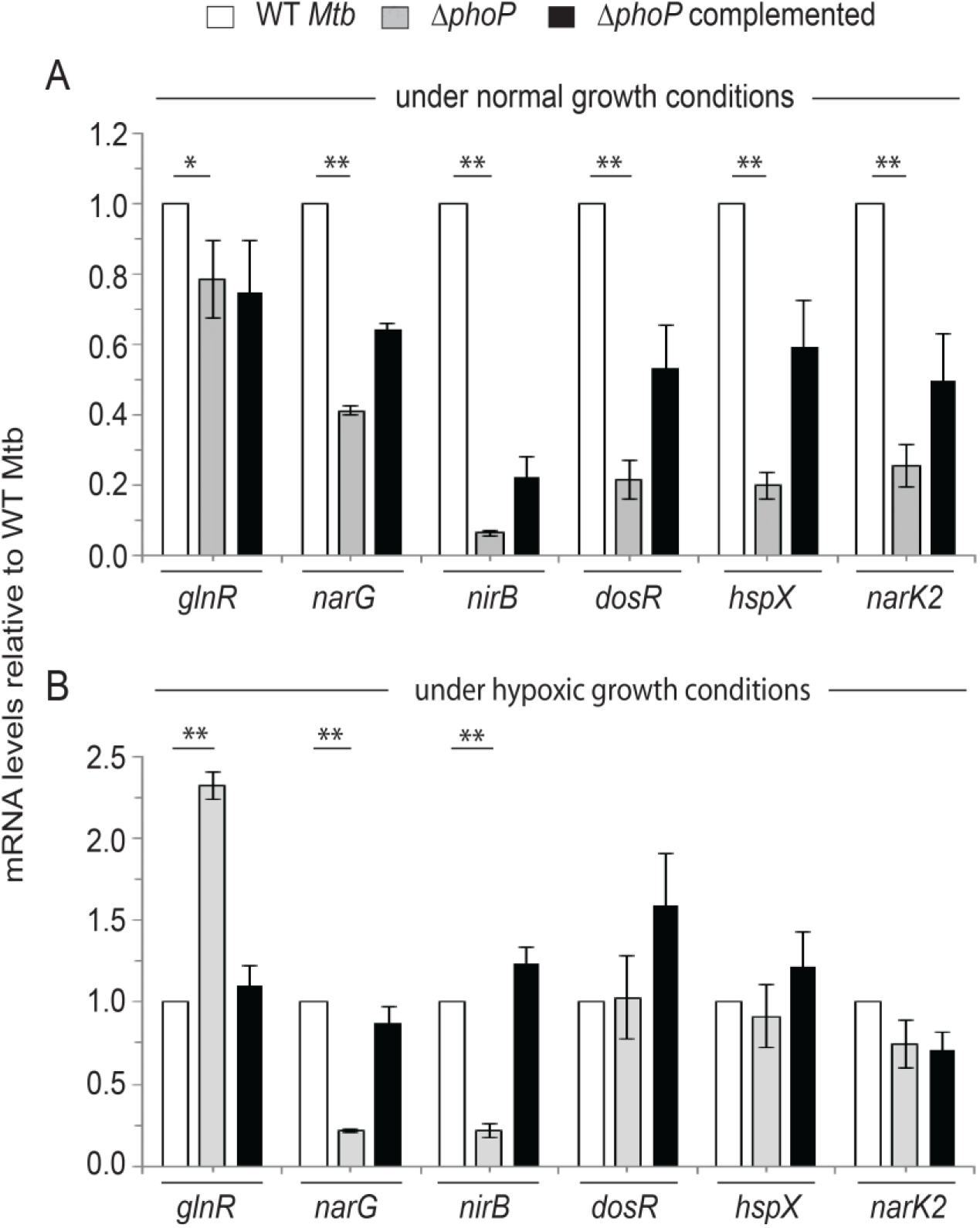
Mtb *phoP* regulates expression of hypoxia-responsive genes *in vivo*. Real-time RT-qPCR was carried out to determine relative expression levels of indicated hypoxia-inducible genes in WT (wild-type), Δ*phoP* and the complemented mutant both (A) under normal conditions and (B) hypoxia, as described in the Materials and Methods. The fold difference of mRNA levels with standard deviations from replicate experiments were determined from at least three independent RNA preparations (\*\**P*<0.01).

Next, to investigate expression levels of *glnR*, *narG* and *nirB*, we grew WT Mtb in Dubos medium with either limiting (1 mM) or surplus nitrogen (30 mM) conditions (Fig. S1), as described earlier (32). Although mycobacterial genes involved in N_2_ metabolism showed a relatively lower level of expression under hypoxia coupled with limiting ammonium chloride concentration (Fig. S1A), these genes displayed a significant induction during hypoxia coupled with surplus nitrogen (Fig. S1B). In sharp contrast, regardless of the ammonium chloride concentration, mycobacterial genes *dosR*, *hspX* and *narK2* showed a significantly higher expression level under normoxia (compare Fig. S1A and Fig. S1B).

To examine whether PhoP controls expression of mycobacterial N_2_ metabolism, we grew WT and *ΔphoP* Mtb in Dubos medium with either limiting nitrogen (1 mM) or nitrogen surplus (30 mM) conditions (Fig. 2). In agreement with the above results (Fig. 1), during hypoxia coupled with surplus nitrogen concentration, expression of *glnR*, *narG* and *nirB* was dependent on PhoP (Fig. 2A). In contrast, (a) under identical conditions, *phoP* locus had no effect on expression of *dosR*, *hspX* and *narK2* (Fig. S2A), and (b) under normoxia coupled with surplus ammonium chloride concentration, these genes showed comparable expressions in WT and *ΔphoP* mutant (Fig. 2B). On the other hand, regardless of the ammonium concentration, under normoxia *phoP* locus shows a striking effect on the expression of hypoxia-responsive genes *dosR*, *hspX*, and *narK2* (Fig. S2B). From these results, we conclude that *phoP* plays a major regulatory role in mycobacterial nitrogen metabolism under oxygen-limiting (hypoxia) conditions. A regulatory scheme suggesting activation of *narG* and *nirB* and repression of *glnR* to lower nitrogen scavenging is shown in Fig. 2C.

**Fig. 2:**
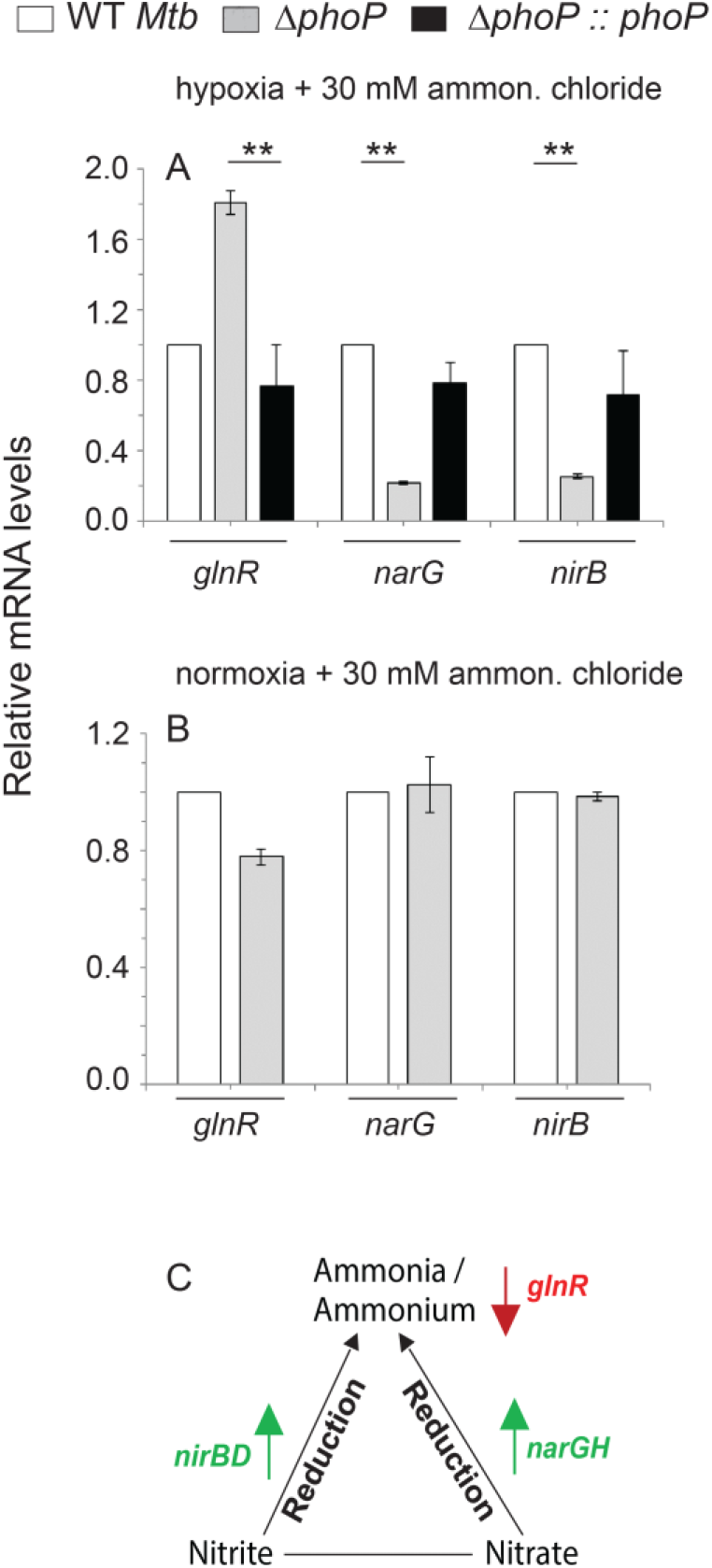
Regulation of genes related to nitrogen metabolism by the *phoP* locus. (A-B) RT-qPCR was carried out to compare expression levels of hypoxia-inducible genes in the WT, *ΔphoP* and the complemented mutant under specific and indicated conditions of growth. The difference of mRNA levels with standard deviations were determined from at least three independent RNA preparations (\*\**P*<0.01). (C) The scheme summarizes *phoP* dependent regulation of nitrogen metabolism during hypoxia. While PhoP acts as an activator of nitrite and nitrate reductases (*nirB* and *narG*, respectively) during hypoxia (as shown by upward green arrows), repression of *glnR* (as shown by a downward red arrow) signifies lower scavenging of nitrogen source under oxygen austerity.

### Regulation of hypoxia-responsive promoters by PhoP and DosR

To examine PhoP dependent *dosR* expression, we probed PhoP binding site within the *dosR* promoter (dosRup spanning −1438 to −774 relative to the start of the ORF). Based on the knowledge of consensus PhoP binding site (31, 32), and more recent results on PhoP binding (23) we could locate a region spanning −1117 to −1100 within dosRup as the likely PhoP box. Next, mutant dosRup (dosRupmut) was generated by introducing mutations at the PhoP box as shown in Fig.3A, and transcriptional fusions of the WT (dosRup) and the mutant promoter (dsoRupmut) were cloned at the ScaI site of pSM128 (33), a promoter-less integrative lacZ reporter vector with a streptomycin resistance gene. Mtb PhoP was expressed in *M. smegmatis* by using pME1mL1-*phoP* as described previously (34) (see Methods for details). Consistent with *phoP*-dependent activation of *dosR* expression (Fig. 1), the dosRup-*lacZ* fusion showed a significant activation (4±0.9-fold) of promoter activity with induction of *phoP* expression relative to the un-induced culture (Fig. 3B). However, dosRupmut -*lacZ* showed comparable β-galactosidase activity with or without induction of PhoP (1.2±0.05-fold difference). Inset to Fig. 3B compares Mtb PhoP expression in *M. smegmatis* harboring WT or the mutant promoter. These results establish that PhoP-dependent activation of *dosR* expression involves the above-noted direct repeat motif as the primary target site of PhoP.

**Fig. 3:**
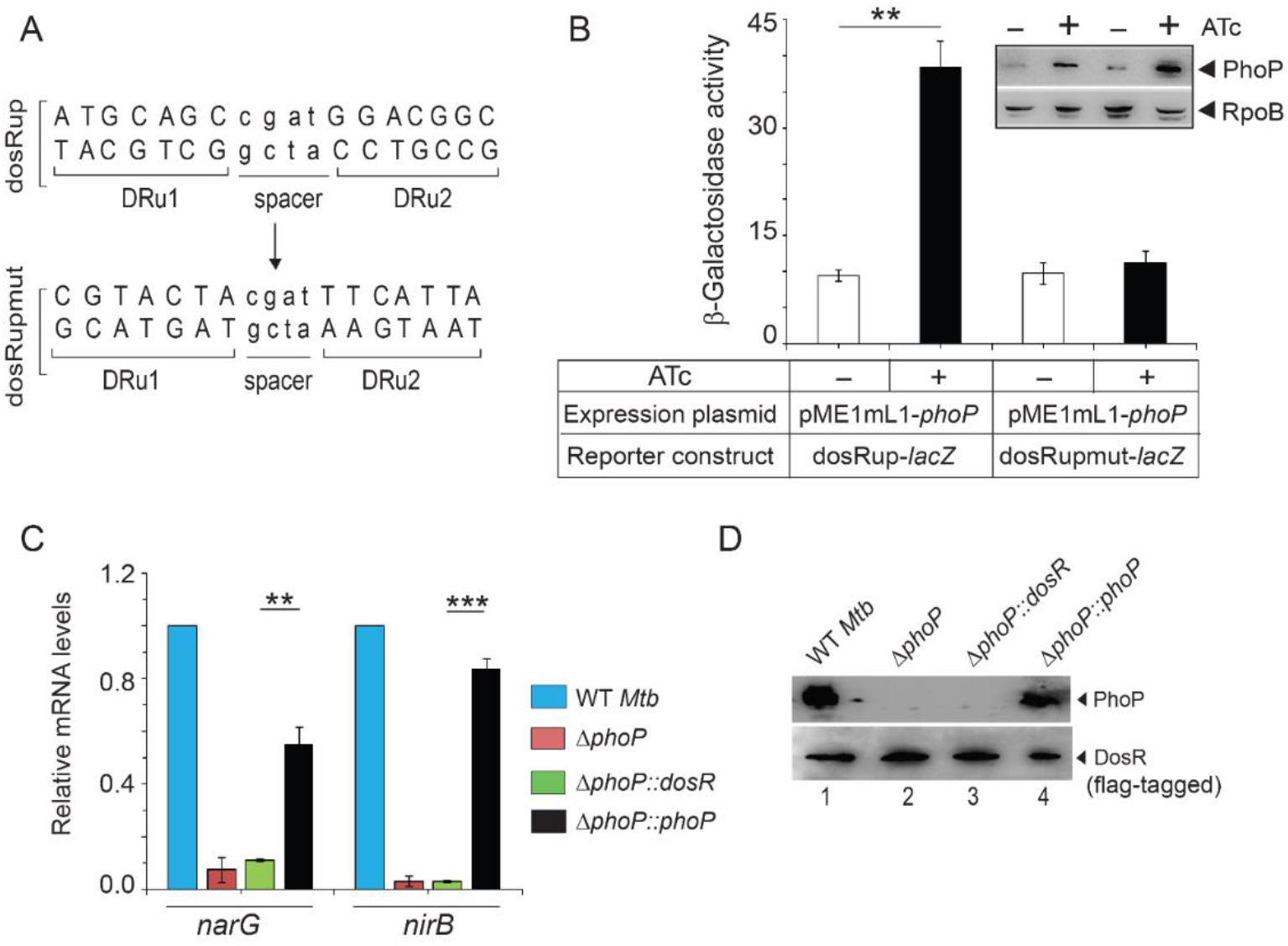
Regulatory mechanism of hypoxia-inducible promoter activity by PhoP. (A) PhoP binding direct repeat motif showing upstream (DRu1) and downstream (DRu2) repeat units. The mutant promoter of *dosR* carrying changes only at the PhoP binding site, dosRupmut, was generated by changing As to Gs and Cs to Ts (of both repeat units) and *vice versa*. Next, the orientation of the DRu2 sequence was reversed relative to DRu1. (B) *M. smegmatis* strains harboring either WT (dosRup-*lacZ*) or the mutant promoter (dosRupmut-*lacZ*) construct along with PhoP expression plasmid (pME1mL1-*phoP*) were grown for 24 hours in absence (−) or presence (+) of ATc, as inducer of PhoP expression. β-galactosidase activities were determined as described in the methods section. The activities represent average of multiple experiments with standard deviations from at least three independent cultures. Inset compares expression of PhoP in ~10 μg crude extracts as detected by anti-PhoP antibody; RpoB was used as the loading control. (C) FLAG-tagged DosR was cloned and expressed in mycobacterial expression vector p19Kpro (51) as described in the methods section. To examine effect of *dosR* expression in *ΔphoP*, mRNA levels of hypoxia-responsive *narG* and *nirB* were determined in indicated strains by RT-qPCR analyses, as described in the legend to Fig. 1. (D) To verify expression of PhoP and DosR in indicated strains, mycobacterial cell extracts containing comparable amount of total proteins were resolved by SDS-PAGE, and the presence of the regulators were confirmed by Western blot analyses using anti-PhoP and anti-FLAG antibodies, respectively.

To examine whether *phoP*-dependent regulation of hypoxic response is achieved via DosR expression, we next expressed *dosR* in *ΔphoP* Mtb and compared expression levels of *narG* and *nirB* (Fig. 3C). Our results unambiguously show that *ΔphoP∷dosR* was unable to complement *narG* and *nirB* expression relative to *ΔphoP∷phoP*, suggesting that *phoP*-controlled expression of *narG* and *nirB* is not attributable to *phoP*-dependent activation of *dosR* expression. Fig. 3D confirms expression of regulators in indicated strains. These results suggest that ectopic expression of *dosR* in *ΔphoP∷dosR* is unable to complement *narG* and *nirB* expression. However, complementation of *ΔphoP* under identical conditions could complement target gene expressions. From these results, we conclude that PhoP-dependent *dosR* expression cannot account for its regulation of hypoxic response. This observation fits well with the above finding that PhoP-dependent regulation of hypoxia-inducible genes (*dosR*, *hspX* and *narK2*) is limited to normoxia only (Fig. 1), and thus possibly remain unlinked to hypoxic response of Mtb

As both PhoP and DosR was shown to regulate hypoxia-responsive genes, we next compared *in vivo* recruitment of DosR within hypoxia-responsive promoters of WT and *ΔphoP* (Fig. 4A). This is because hypoxia-responsive *narG*, *nirB* and *glnR* showed a *phoP*-dependent expression during hypoxia (Fig. 1). Note that despite a comparable level of DosR in the WT and *ΔphoP* under hypoxia (inset to Fig. 4A), we observed a significantly lowered DosR recruitment (2.2-6.2 ±0.15 -fold) within the target promoters in *ΔphoP* relative to the WT bacilli. Thus, presence of PhoP is necessary for effective recruitment of DosR within the hypoxia-inducible promoters involved in mycobacterial N_2_ metabolism. Notably, this finding is consistent with previously-reported *dosR* dependent regulation of *narG* expression (13).

**Fig. 4:**
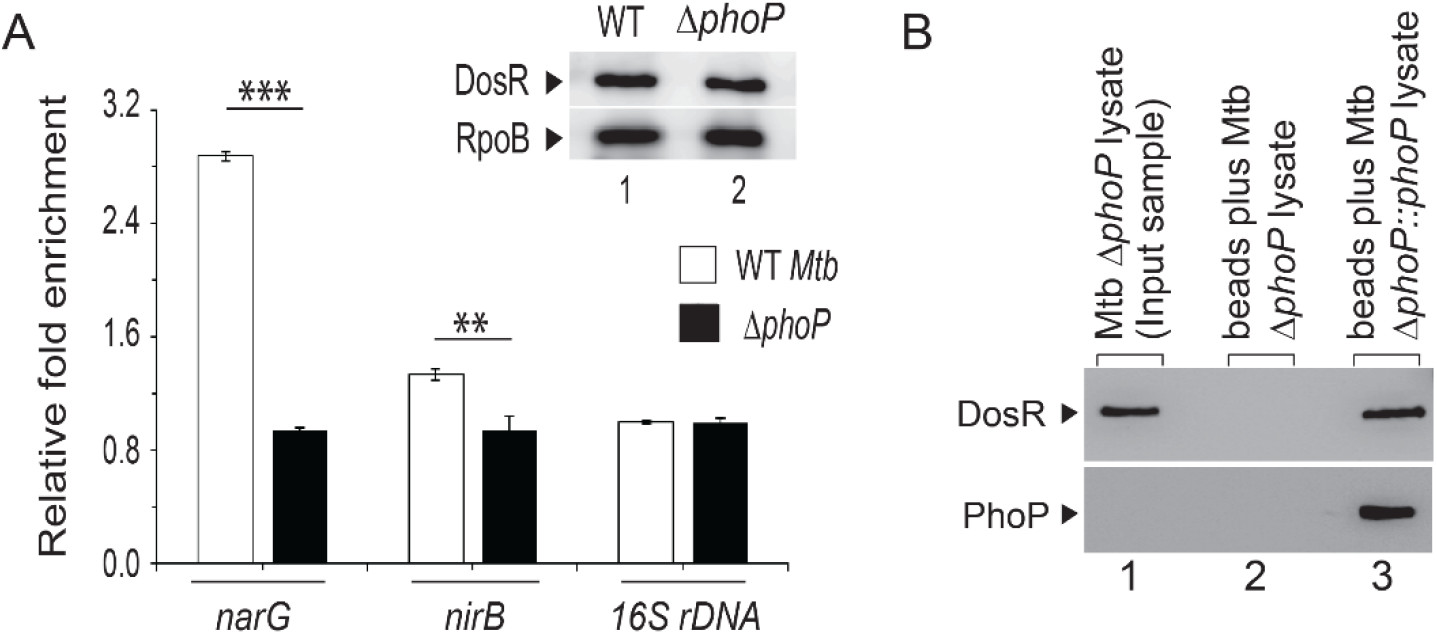
Recruitment of PhoP and DosR within hypoxia- inducible promoters. (A) *In vivo* experiments compared recruitment of DosR in WT and *ΔphoP* grown under hypoxia. ChIP-qPCR experiments utilized anti-FLAG antibodies (Thermo Scientific) to determine fold enrichments with respect to mock IP (without adding antibody) sample, as described in the ‘methods’ section and inset compares DosR expression in 10 μg crude cell-lysates; RpoB was used as the loading control. (\*\**P*-value <0.01; \**P*-value<0.05). (B) Crude cell lysates of *ΔphoP* expressing His_6_-tagged PhoP (p19kpro-*phoP*; Table S3) was incubated with Ni-NTA and eluted with 200 mM imidazole; lane 1, input sample; lane 2, elution from the crude lysate of cells lacking *phoP* expression; lane 3, detecting DosR co-elution with PhoP; upper and the lower panels identify DosR and PhoP by anti-DosR and anti-His antibody, respectively using Western blotting as described in the Methods.

Considering the above results, recently-reported PhoP-DosR interaction (23) emerged as a possible explanation to account for the regulation of hypoxia-inducible genes. To examine this, whole cell lysate of *ΔphoP* expressing His-tagged PhoP, as described previously (35), was incubated with Ni-NTA, and proteins were eluted from Ni-NTA column using imidazole (Fig. 4B). While the eluent showed clear presence of DosR (lane 3), we were unable to detect DosR from the cell lysate of Δ*phoP* carrying the empty vector (p19Kpro) (lane 2, see Table S3). In agreement with recently reported PhoP-DosR interaction by Vashist et al. (23), these results suggest specific *in vivo* interaction between PhoP and DosR.

Next, we utilized mycobacterial protein fragment complementation (M-PFC) assay to examine PhoP-DosR interaction using *M. smegmatis* as the surrogate host (Fig. S3) (see Table S3 for plasmids used in this study). In agreement with results reported previously (23), our results also showed specific interaction between PhoP and DosR (Fig. S3A). However, under identical conditions, we were unable to detect interaction between PhoP and GlnR (Fig. S3B), suggesting that regulation of *narG* and *nirB* expression during hypoxia is most likely GlnR independent. Using phosphorylation defective mutants of PhoP and DosR (PhoPD71N and DosRD54N, respectively), we further showed that phosphorylation of the response regulators do not appear to influence PhoP-DosR protein-protein contacts (Figs. S3C-D).

### Probing PhoP-DosR protein-protein interactions

To investigate interacting domains of PhoP and DosR, we next performed *in vitro* protein-protein interaction analysis using GST-DosR and domains of His_6_-tagged PhoP (Fig. 5A), previously shown to be functioning on their own (36). In pull down assays, GST-DosR was immobilized on glutathione-sepharose followed by incubation with His_6_-tagged PhoP domains (Fig. 5A). Upon elution PhoPN (comprising PhoP residues 1-141; lane 1) is co-eluted with GST-DosR from the bound glutathione sepharose by 10 mM glutathione [similar to PhoP (lane 4)]. However, under identical conditions, none of the PhoP C-domain constructs (PhoPC1, comprising residues 141-247 or PhoPC2, comprising residues 150-247) is co-eluted with GST-DosR (lanes 2- 3).

**Fig. 5:**
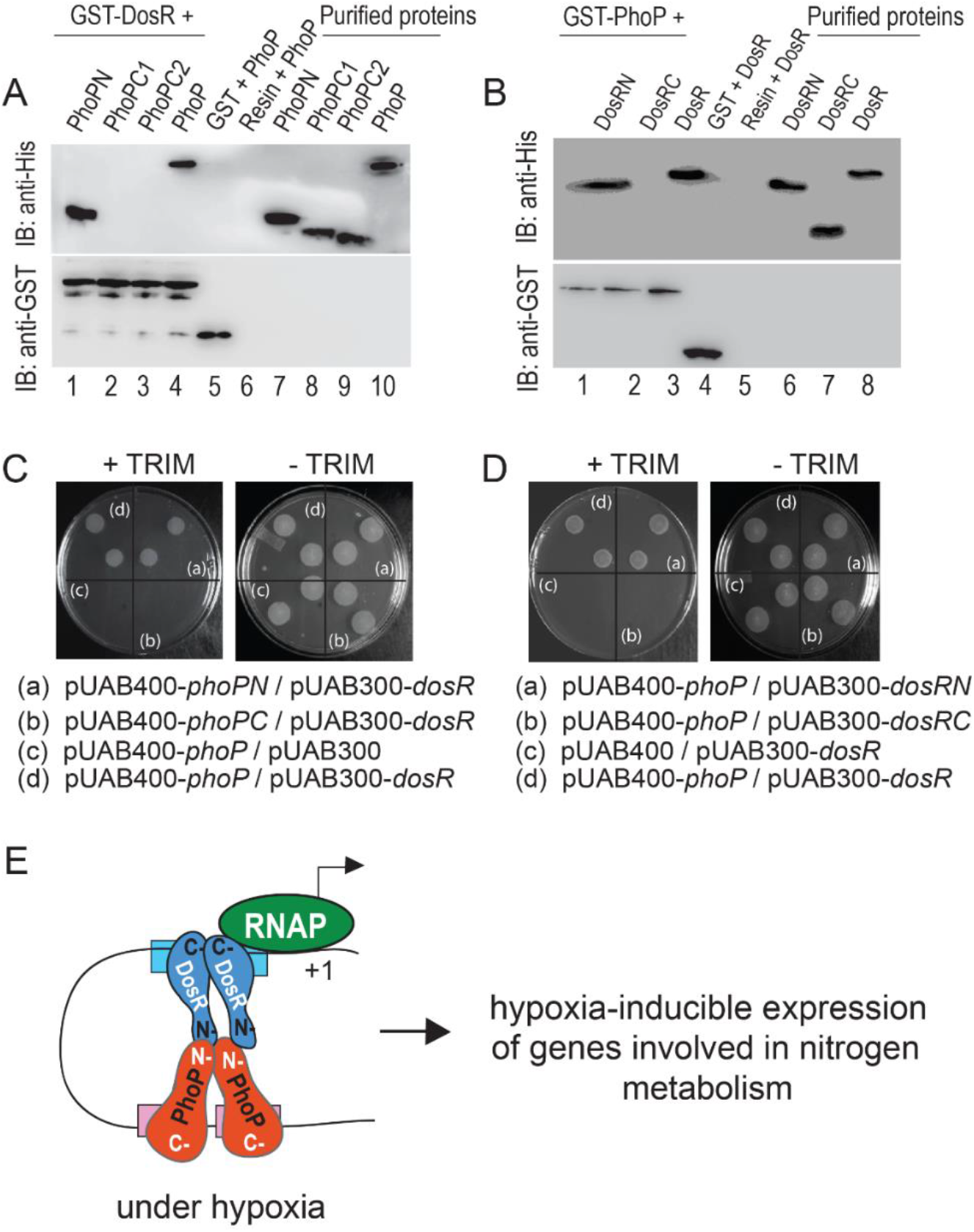
Probing interacting domains of PhoP and DosR. (A) Indicated His_6_-tagged PhoP domains were incubated with glutathione sepharose, previously immobilized with GST-DosR. Fractions of bound proteins (lanes 1-4) were analyzed by anti-His (upper panel) or anti-GST antibody (lower panel). Control sets include GST alone (lane 5), or the resin alone (lane 6); lanes 7-10 resolve purified PhoP constructs. (B) Likewise, indicated His_6_-tagged DosR domains were incubated with glutathione sepharose previously immobilized with GST-PhoP. Fractions of bound proteins (lanes 1-3) were probed with anti-His or anti-GST antibody (as above). Control sets include GST alone (lane 4), or the resin alone (lane 5); lanes 6-8 resolved purified DosR constructs. The results suggest that the DosRN, and not DosRC, retains the ability to interact with PhoP. (C-D) M-PFC experiments examined interactions between (C) indicated PhoP domains and DosR, or (D) PhoP and indicated DosR domains, respectively, using full-length PhoP and DosR pair as the positive control. (E) Model depicting how PhoP and DosR function as co-activators of hypoxia-inducible gene expression (right).While PhoP-DosR interaction via their received domains contribute to additional stability of the transcription initiation complex, mycobacterial RNA polymerase bind to the target site of the promoter to initiate transcription.

To identify the corresponding interacting domain of DosR, in similar pull-down assays as above, we used GST-PhoP and His-tagged DosR-domain constructs (Fig. 5B, see Table S3). Interestingly, DosRN (comprising residues 1-193) co-eluted with GST-PhoP (lane 1); however, DosRC (comprising residues 143-217) under identical conditions, did not co-elute with GST-PhoP (lane 2), suggesting specific interaction between DosRN and PhoP. Fig. S4 shows the purified form of the recombinant regulators (and their domains) used in these experiments. Consistent with the above results, *M. smegmatis* cells co-expressing PhoPN and DosR (Fig. 5C) as well as PhoP and DosRN (Fig. 5D) pairs grew well on TRIM plates. However, cells co-expressing PhoPC and DosR or PhoP and DosRC pairs failed to grow on TRIM plates (compare Figs. 5C, and 5D). Note that all of these cells grew well on plates lacking TRIM. Together, these results suggest that N-domain of PhoP interacts with the N-domain of DosR. Based on the promoter occupancy of PhoP (22, 31) and DosR (15, 37) and the orientation of the two interacting regulators, Fig. 5E summarizes a regulatory scheme showing co-activation of hypoxia-inducible genes (involved in N_2_ metabolism) by PhoP and DosR.

### phoP -dependent growth restoration of Mtb under hypoxia

With the results showing a striking impact of PhoP on expression of hypoxia-inducible genes involved in N_2_-metabolism, we sought to examine growth of WT and *ΔphoP* under hypoxia (Fig. 6). When compared under hypoxia coupled with N_2_-limiting conditions, we observed that both strains showed comparably limited growth (Fig. 6A). However, under hypoxia coupled with N_2_-surplus conditions, WT bacilli showed significant growth restoration (Fig. 6B). In striking contrast, *ΔphoP* Mtb under identical conditions, failed to resume growth as that of the WT bacilli, and consistently showed a lower growth of 2±0.1 fold relative to the WT bacilli. Importantly, growth defect of *ΔphoP* could be restored to the WT level with the complementation of the mutant (see also Fig. S5), underscoring the importance of *phoP* in utilizing available nitrogen resource during hypoxia. As controls, under normoxia, WT and *ΔphoP* grew comparably well regardless of available ammonium chloride concentration (Figs. 6C-D). Together, we conclude that under hypoxic conditions, PhoP integrates nitrogen metabolism and hypoxia.

**Fig. 6:**
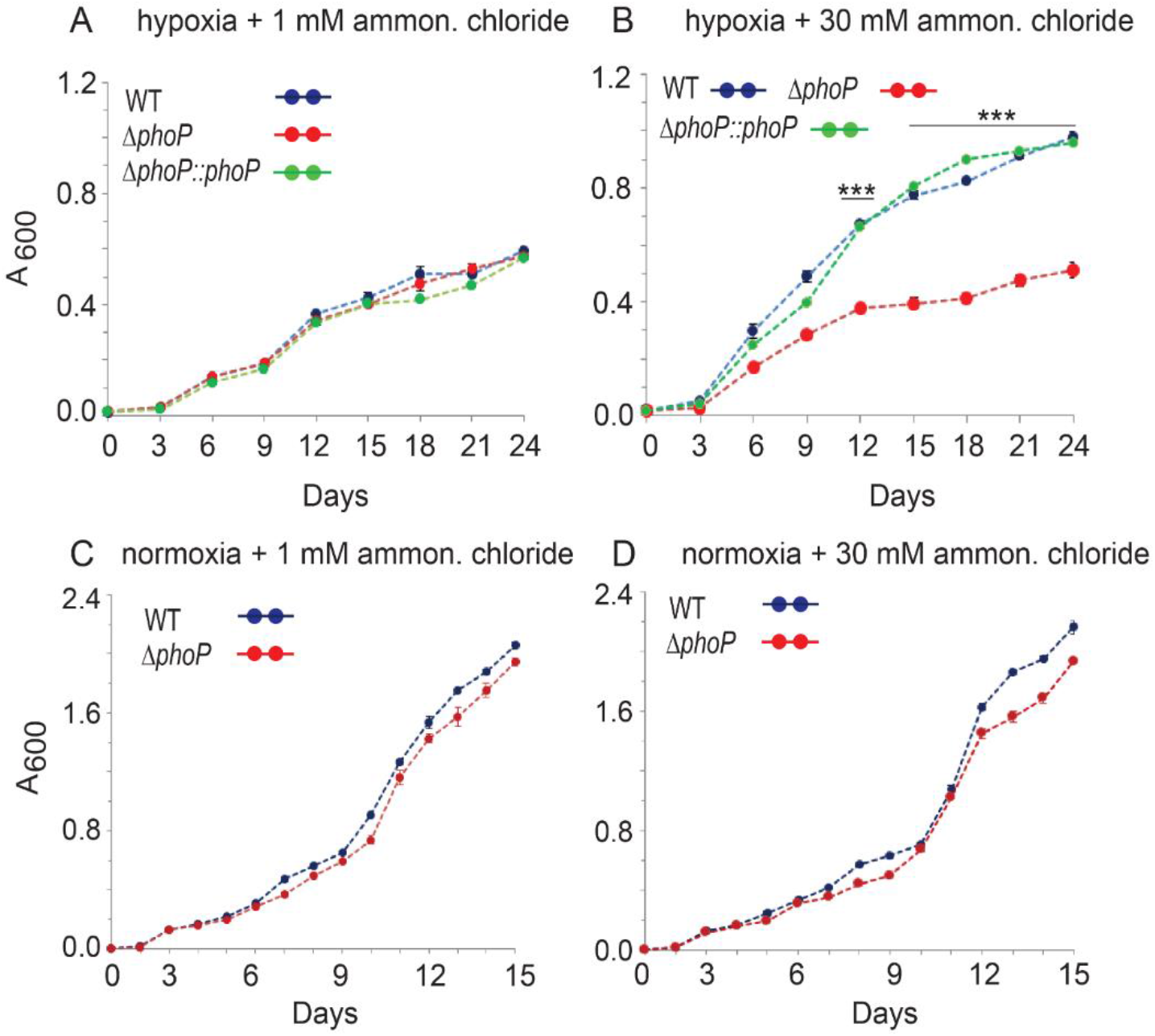
Mycobacterial growth under hypoxia coupled with surplus or limiting nitrogen availability. (A-D) Growth curves of indicated strains were compared under hypoxia (A-B) or normoxia (C-D) coupled with either limiting nitrogen (1 mM NH4Cl) or surplus nitrogen (30 mM NH4Cl) conditions. The values represent average of replicate experiments with standard deviations from at least three independent cultures (\*\*\**P*<0.001).

## DISCUSSION

Mtb, during infection survive in environments with oxygen limiting conditions, yet the mechanisms that promote survival of the metabolically inactive bacilli remains largely unknown. The fact that oxygen pressure within granulomas remains low (38, 39), yet reactivation from latency takes place within oxygen rich sites of the lung, suggest that oxygen availability remains the key determinant. In keeping with this, mycobacterial growth both *in vitro* and *in vivo*, are strongly influenced by the available oxygen pressure (2, 40), suggesting hypoxia as one of the major trigger factors of latency and reactivation.

Although hypoxia is known to increase nitrate reduction in Mtb (7), mechanism of regulation of nitrogen metabolism during oxygen austerity remained obscure. While DosR remains essential for Mtb adaptation and survival under hypoxic environment, PhoP coordinates pH homeostasis via regulation of pH-inducible gene expression. Interestingly, DosR was recently implicated in Mtb growth at low pH under anaerobic conditions (24). Thus, we sought to investigate role of PhoP on hypoxic response of Mtb. Results reported in this study suggest that PhoP activates expression of mycobacterial genes related to N_2_ metabolism under oxygen limiting conditions with effective assistance from the hypoxia regulator DosR. Together, our results provide a new biological insight showing metabolic switching of Mtb in response to hypoxia by the convergence of two major response regulators.

It is noteworthy that Mtb, which evolved as an obligate human pathogen, has lost majority of the regulatory pathways that are part of metabolism of easily available nutrients. Thus, nitrogen-metabolism regulon is largely restricted to nitrite and nitrate reductases, making available the most likely nitrogen sources within the host (41). Notably, nitrate remains the final electron acceptor in lieu of oxygen to support growth of many bacteria during oxygen austerity. Therefore, hypoxia-induced non-replicating persistence in Mtb is accompanied with nitrate reduction as a way to maintain redox-balance and save energy reservoir during shift-down (8). In keeping with these results, under hypoxia, (a) activation of *nirBD* and *narGHIJ* loci, which function in nitrite and nitrate reductions, respectively, and (b) repression of *glnR*, which controls nitrogen scavenging (41) by the *phoP* locus unravel a strikingly novel role of PhoP in nitrogen metabolism during hypoxia.

Considering the involvement of both the regulators, it was of interest to examine whether these two are functionally connected. Clearly, recruitment of PhoP and DosR (Fig. 4A-B), most likely controlled by protein-protein interactions (Fig. 4), regulate activation of hypoxia-inducible genes, provides an integrated view of our results. Under such a situation, for a more effective functioning DosR recruitment is possibly ensured within target promoters already bound by PhoP. Based on the involvement of both regulators, coupled with our findings that the respective N-domains interact to each other (Figs. 5A-D), we propose a model (Fig. 5E) suggesting how PhoP-DosR interaction controls precise regulation of hypoxic response, a key step in the intracellular survival of Mtb. These considerations take on more significance in the light of previously-identified PhoP and DosR binding sites within the afore-noted promoters. The arrangement and spacing between the binding sites is strongly indicative of DNA looping, possibly to assist transcription initiation. Together, these results (a) suggest a critical role of PhoP in binding and transcriptional control of *narG* and *nirB*, and (b) account for an explanation of why DosR was unable to be recruited at these promoters in *ΔphoP* Mtb (Fig. 4B), a finding that fits well with the previously-reported PhoP-DosR interaction data (23) (see Fig. 4). Although the model (Fig. 5E) shows an equimolar PhoP and DosR in the ternary complex, there is no evidence to suggest 1:1 binding stoichiometry. However, as independent regulators, in both cases a dimer of PhoP or DosR has been shown to bind DNA (15, 22, 37). From the ChIP results (Fig. 4), DosR remains ineffective in *ΔphoP* and therefore, *ΔphoP* is expected to display a growth defect under hypoxia. However, we find comparably limited growth by both the WT and the mutant bacteria under hypoxia (Fig. 6A). We argue that PhoP-DosR interaction -dependent regulation possibly controls a few critical genes related to nitrogen metabolism, and therefore a significant growth defect of *ΔphoP* relative to the WT bacilli is apparent only when surplus nitrogen is available during oxygen austerity (Fig. 6B).

Integration of two signaling systems, either similar or of different types are known in numerous other biological systems. An earlier study demonstrates role of PknH in phosphorylation of DosR, necessary for complete induction of the Mtb *dosR* regulon (42). While these results suggest convergence of two different types of signalling modules for a common function, along the similar line DosR interacts with the house keeping sigma factor SigA for bacterial survival under dormancy (17). Recently, we have shown that PhoP interacts with nucleoid-associated protein EspR to control ESAT-6 secretion (35). In this study, we show that N-domains which get phosphorylated for activation of respective regulators, interact to each other (Fig. 5). This is in sharp contrast to recently-reported PhoP-HspR and PhoP-HrcA interactions where PhoPN was shown to interact with the C-terminal end of the mycobacterial heat-shock repressors (43). Although phosphorylation of either of the regulators does not seem to influence PhoP-DosR protein-protein contacts (Fig. S3C & D), involvement of additional regulatory control by the N-domains other than phosphorylation (Fig. 5), which enables appropriate *in vivo* functioning of the regulators via protein-protein contacts (Fig. 4), offers a new mechanistic insight. However, from these results we cannot rule out involvement of PhoR and DosS/DosT during hypoxia. In fact, a previous study suggests cross-talk between the two signalling systems (44). As several examples establish that either two different or similar regulatory systems are often integrated toward a common regulatory function, integration via a single interaction (or lack thereof) possibly could have a large or a small impact on transcriptional control of different target promoters. It is conceivable that diversity of interactions at numerous promoters (belonging to various regulons) may significantly enhance the available combinations of potential regulatory interactions to fine tune context-dependent gene expression. As hypoxic response is believed to play a major role in dormancy adaptation, DosR activation is associated with metabolic changes where the tubercle bacilli move into a non-replicating persistent state. Therefore, it is not too difficult to imagine that control mechanisms (for example, PhoP-DosR interactions) would exist to interfere with bacterial persistence unless an appropriate signal is recognized. In fact, this is expected since activation of the regulon consisting of ~48 genes would otherwise be energy-demanding, and therefore, it appears that a second molecular control system (in addition to DosS/DosT) would be in place to regulate induction of DosR only under the most appropriate stress conditions.

In conclusion, we have identified a signaling mechanism in Mtb that regulates hypoxia-inducible genes related to nitrogen metabolism. To our knowledge, this is the first report of a regulatory pathway that links the virulence regulator PhoP to expression of nitrite and nitrate reductases under hypoxia. Together, this study explains, at least in part, a fundamental mechanism of metabolic switching underlying how nitrogen metabolism genes are activated for survival of Mtb under oxygen austerity for a long period of time.

## Materials and Methods

### Bacterial strains and culture conditions

*E. coli* DH5α, and *E. coli* BL21 (DE3) strains were grown at 37°C in LB media containing appropriate antibiotics, and used for cloning and for over-expression of mycobacterial proteins, respectively. While *ΔphoP* and the complemented mutant have been described previously (45), *ΔdosR* and the complemented mutant are described below. Mtb strains were grown aerobically at 37°C in Middlebrook 7H9 liquid broth (containing 0.2% glycerol, 0.05% Tween-80 and 10% ADC) or on 7H10-agar medium (containing 0.5% glycerol and 10% OADC). For growth under hypoxic conditions, stock cultures were aerobically sub-cultured twice to mid-log phase (A_600_ of 0.3-0.4) in Dubos (Difco) media supplemented with 0.045 % Tween 80, 10 % albumin and dextrose, and subsequently inoculated in fresh media at A_600_ of 0.01. Oxyrase was used to remove excess of oxygen and 1.5 μg/ml methylene blue was added as an indicator of oxygen depletion (7). Mtb growth under nitrogen excess or nitrogen limiting conditions are as described (41), and the expression of *nirB* was measured by RT-qPCR using RNA from three independent cultures with *gapdh* as internal control. Briefly, indicated Mtb strains were grown as above, washed twice in nitrogen-free Dubo’s medium, and inoculated in the same medium either supplemented with 1 mM (nitrogen limiting) or 30 mM (nitrogen surplus) ammonium chloride (Sigma). Transformation of wild-type (WT) and mutant Mtb strains and selection of transformants on appropriate antibiotics were carried out as described previously.

### RNA isolation and quantitative real-time RT-PCR

Total RNA from Mtb was extracted as described (28) using exponentially growing bacterial cultures grown with or without stress. After extraction with chloroform-isoamyl alcohol, RNA was precipitated with chilled ethanol. Finally RNA was treated with RNase-free DNaseI (Invitrogen) for 20 minutes at room temperature to remove genomic DNA contamination. The integrity of RNA samples was checked by intactness of 23S and 16S rRNA using formaldehyde-agarose gel electrophoresis. RNA concentrations were determined by measuring absorbance at 260 nm, and stored at −80°C.

cDNA synthesis and PCR reactions using appropriate PAGE-purified primer pairs (200 nM; see Table S1) were performed in an Applied Biosystems real-time PCR detection system using Superscript III platinum-SYBR green one-step qRT-PCR kit (Invitrogen) and cycling conditions as described (28). For each pair of primers, a standard curve was generated using serially diluted RNA samples, and PCR efficiency was evaluated. The average fold change of expression of each sample relative to endogenously expressed Mtb *gapdh* (Rv1436) was calculated by the ΔΔC_T_ method (46). Note that CT values for *gapdh* remained mostly unchanged under variable conditions of Mtb growth and standard deviations were derived from at least three independent RNA preparations. Platinum Taq DNA polymerase (Invitrogen) was used to confirm absence of genomic DNA in our RNA preparations.

### Cloning

Isolation and purification of nucleic acids, digestion with restriction enzymes, and analyses of nucleic acids or its fragments by agarose gel electrophoresis followed standard procedures. Mtb PhoP or its domain-specific over-expressing constructs have been described earlier (47). Likewise, full length DosR (encoding 654-bp) ORF, and truncated DosR proteins (encoding N-terminal 579-bp, and C-terminal 222-bp of the *dosR* ORF) were cloned in T7-lac-based expression system pET28b (Novagen) as recombinant fusion proteins containing an N-terminal His_6_-tag. The cloning strategy resulted in pET-*dosR*, pET-*dosRN*, and pET-*dosRC* comprising 217 amino acid full-length DosR, 193 amino acid DosRN (lacking the C-terminal 24 residues), and 74 residues long DosRC (lacking the N-terminal 142 residues), respectively (48). Plasmid pGEX-*dosR* expressing DosR with an N-terminal GST-tag was generated by cloning PCR-derived *dosR* ORF fragment between BamHI and XhoI sites of pGEX 4T-1 (GE Healthcare) as described for pGEX-*phoP* (49). To complement *dosR* expression in the respective mycobacterial mutant, the ORF was cloned and expressed in mycobacterial expression vector pSTKi (50). To express FLAG-tagged *dosR* in Mtb, the ORFs were cloned and expressed in mycobacterial expression vector p19Kpro (51). Point mutation in *phoP* and *dosR* genes were introduced by two-stage overlap extension method, and verified by DNA sequencing. The oligonucleotide primers used in cloning and construction of the plasmids are provided in Table S2, and Table S3, respectively.

### Promoter regulation by Mtb PhoP in *M. smegmatis*

*M. smegmatis* strains carrying indicated *lacZ* fusions and pME1mL1-*phoP*, expressing Mtb PhoP under P_myc1_*tet*O promoter and TetR repressor (or no expression plasmid as control), were grown either in absence or in presence of 50 ng/ml of anhydrotetracycline (ATc) as described previously (52). To determine promoter activity, cells from both induced and un-induced cultures were grown for 24 hours, cell suspensions were sonicated, and β-galactosidase activity of the extracts were determined. To assess TetR-dependent PhoP expression, crude cell lysates of *M. smegmatis* (≈10 μg protein) were resolved by 12% SDS-PAGE and probed with anti-PhoP antibody (AlphaOmega Sciences, India).

### Proteins

Mtb PhoP and its domains were expressed and purified as described (36). Full-length and truncated DosR proteins were expressed in *E. coli* BL21 (DE3) as fusion proteins containing an N-terminal His_6_- tag and purified by immobilized metal-affinity chromatography (Ni-NTA, Qiagen). Full-length DosR was also expressed with N-terminal GST-tag, as described for GST-PhoP (53). Finally, the proteins were stored in buffer containing 50 mM Tris-HCl, pH 7.9, 300 mM NaCl, and 10% glycerol. In all cases the purity was verified by SDS-PAGE, protein concentrations were determined by Bradford reagent with BSA as the standard, and expressed in equivalent of protein monomers.

### Immunoblotting

Crude cell lysates of Mtb were resolved by 12% SDS-PAGE and visualized by Western blot analysis. For immunoblotting, resolved samples were electroblotted onto polyvinyl difluoride (PVDF) membranes (Millipore) and were detected by affinity-purified antibodies elicited in rabbit (AlphaOmega Sciences). RNA polymerase, used as a loading control, was detected by anti-RpoB (Abcam). Anti-His and anti-GST antibodies were from GE Helathcare. Goat anti-rabbit and goat anti-mouse secondary antibodies conjugated to horseradish peroxidase were procured from AlphaOmegaSciences, and blots were developed with Luminata Forte chemiluminescence reagent (Millipore).

### ChIP-qPCR

Mtb was grown as described above and processed for ChIP experiments using purified anti-PhoP (Alpha Omega Sciences) or anti-FLAG (Thermo Scientific) antibody and protein A/G agarose beads (Pierce) as described (54). qPCR was performed using PAGE purified primer pairs (Sigma; Table S1) that spanned appropriate promoter regions of interest. PCR reaction mix contained suitable dilutions of immunoprecipitated (IP) DNA in a reaction buffer containing SYBR green mix, and specific primers (200 nM). An IP experiment without adding antibody (mock) served as a negative control, and data was normalized against PCR signal from mock sample. Typically, 40 cycles of amplification were carried out using an Applied Biosystems real-time PCR detection system with serially diluted DNA samples (mock, IP treated and total input). Melting curve analysis was carried out to confirm amplification of a single product in all cases. Enrichment of PCR signal from anti-PhoP or anti-FLAG IP relative to the signal from an IP experiment without adding any antibody (mock) was measured to determine efficiency of recruitment. Specific PCR-enrichment was ensured by performing ChIP-qPCR of the identical IP samples using *gapdh/16S rDNA* specific primers. Each data point represents the mean of duplicate qPCR measurements using at least three independent Mtb cultures.

### Mycobaterial protein fragment complementation (M-PFC) assays

To express Mtb PhoP or its domains in *M. smegmatis*, *phoP* and/or its appropriate domain constructs were cloned in the integrative vector pUAB400 (kan^R^; Table S3) between MfeI and HindIII sites, as described (53). Similarly, *dosR*, *glnR* and truncated *dosR* ORFs were cloned in episomal plasmid pUAB300 (hyg^R^; Table S3) between BamHI/HindIII sites to generate pUAB300-*dosR*, pUAB-*glnR*, pUAB300-*dosRN*, and pUAB300-*dosRC*, respectively. Next, co-transformed cells were selected on 7H10/Kan/Hyg plates in absence or presence of 15 μg/ml TRIM (Trimethoprim, Sigma) as described (55). In this assay, two interacting proteins as separate fusion constructs of two domains of murine DHFR when co-expressed in *M. smegmatis*, reconstitute functional mDHFR and enable the bacteria to grow on media containing TRIM. As a positive control, ESAT-6/CFP-10 expressing constructs were used in M-PFC experiments. Note that all of the strains used in this study grew well in absence of TRIM.

## Acknowledgements

We thank Adrie Steyn for pUAB300/pUAB400 plasmids, G. Marcela Rodriguez and Issar Smith for Δ*phoP*, and the complemented mutant strain, Harsh Gaur for purification of recombinant DosR, and Ashwani Kumar for very helpful discussions.

This work was supported by a research grant (to D.S.) from the SERB-Department of Science and Technology (DST) (EMR/2016/004904), Government of India, and by intramural funding from CSIR-IMTECH. P.R.S. and V.A.K. received financial support from DBT and CSIR respectively. The funders had no role in the study design, data collection and interpretation, or the decision to submit the work for publication.

